# Selective attention gates the interactive crossmodal coupling between perceptual systems

**DOI:** 10.1101/207548

**Authors:** Silvia Convento, Md. Shoaibur Rahman, Jeffrey M. Yau

## Abstract

Cortical sensory systems often activate in parallel, even when stimulation is experienced through a single sensory modality [1–3]. Critically, the functional relationship between co-activated cortical systems is unclear: Co-activations may reflect the interactive coupling between information-linked cortical systems or merely parallel but independent sensory processing. Here, we report causal evidence consistent with the hypothesis that human somatosensory cortex (S1), which co-activates with auditory cortex during the processing of vibrations and textures [4–9], interactively couples to cortical systems that support auditory perception. In a series of behavioural experiments, we used transcranial magnetic stimulation (TMS) to probe interactions between the somatosensory and auditory perceptual systems as we manipulated attention state. Acute manipulation of S1 activity using TMS impairs auditory frequency perception when subjects simultaneously attend to auditory and tactile frequency, but not when attention is directed to audition alone. Auditory frequency perception is unaffected by TMS over visual cortex thus confirming the privileged interactions between the somatosensory and auditory systems in temporal frequency processing [10–13]. Our results provide a key demonstration that selective attention can modulate the functional properties of cortical systems thought to support specific sensory modalities. The gating of crossmodal coupling by selective attention may critically support multisensory interactions and feature-specific perception.

## Results and Discussion

Attending to a stimulus feature that can be redundantly signalled through different senses can activate multiple cortical sensory systems. For example, the auditory cortex can be activated by visual stimulation even in the absence of sounds, particularly when attending to stimulus features that are associated with sounds [14,15]. Such coactivation of sensory systems is thought to reflect a number of attention-based operations including the selection, refinement, and binding of sensory representations [16]. Critically, the functional relationship between co-activated cortical systems remains ambiguous. Co-activation could reflect an interactive coupling between the cortical sensory regions. Alternatively, co-activation could merely reflect the parallel, but independent activation of sensory systems. Distinguishing between these possibilities would provide critical insight into the functional role of distributed activity over cortical systems that are traditionally thought to be dedicated to unisensory processing.

Somatosensory cortex (S1) often co-activates with auditory cortex [4,7,9,17]. Coactivation of these sensory cortical systems may reflect the recruitment of cortical networks that support frequency representations encoded by both touch and audition[8,18] and shared neural circuits may underlie the highly-specific perceptual interactions observed between touch and audition in the temporal frequency domain [10,19,20]. If tactile frequency perception recruits cortical systems that also support audition, the connectivity within these networks conceivably fluctuates systematically to enable flexible behaviour. Here, we hypothesized that S1 becomes functionally coupled to cortical systems that support audition specifically when attention is directed to tactile frequency. We used transcranial magnetic stimulation (TMS) to probe interactions between the somatosensory and auditory systems. We wished to infer the dependence of crossmodal coupling on attention state based on systematic behavioural changes associated with TMS manipulation of specific brain regions. In an attentional state predicted to increase coupling within a distributed frequency processing network, we reasoned that TMS-induced activity changes in S1 would propagate through the auditory cortical system and ultimately lead to disrupted auditory frequency perception.

To test our hypothesis, we causally manipulated S1 activity using online TMS while participants performed a discrimination task [10–13] that required them to report which of two stimuli was perceived to be higher in frequency (Fig. 1). Subjects (n = 15) performed the discrimination task in separate blocks that differed according to the sensory modality to which subjects directed their attention. In the unimodal auditory block (Fig. 1a, Unimodal_A_), sounds were presented during both stimulus intervals (auditory-only trials), requiring subjects to attend only to audition. In the unimodal tactile block (Fig. 1a, Unimodal_T_), vibrations were presented during both intervals (touch-only trials), requiring subjects to attend only to touch. In the Mixed blocks (Fig. 1b, Mixed), auditory-only and tactile-only trials were randomly interleaved with crossmodal trials that required subjects to compare the frequency of a sound presented in one interval to the frequency of a vibration presented in the other. Randomization ensured that subjects could not predict stimulus modality during any stimulus interval in the Mixed blocks, thereby forcing subjects to attend to both modalities throughout the blocks. Crucially, by comparing performance on the auditory-only trials, which occurred in both the Unimodal_A_ and Mixed blocks (Fig. 1a,b, asterisks), we tested how manipulating S1 activity using TMS impacted auditory frequency perception under states that differed only with respect to whether subjects also attended to tactile frequency cues. According to our hypothesis, we predicted that TMS over S1 would impair auditory perception only during the Mixed blocks. In our within-subjects design, each subject performed the same behavioural task in two sessions in which they received TMS over S1 or a control site (Fig. 1c). During each session, subjects initially performed the auditory and tactile discrimination task in blocks without TMS (baseline blocks) before completing the Unimodal_A_, Unimodal_T_, and Mixed blocks (Methods).

**Figure 1.**
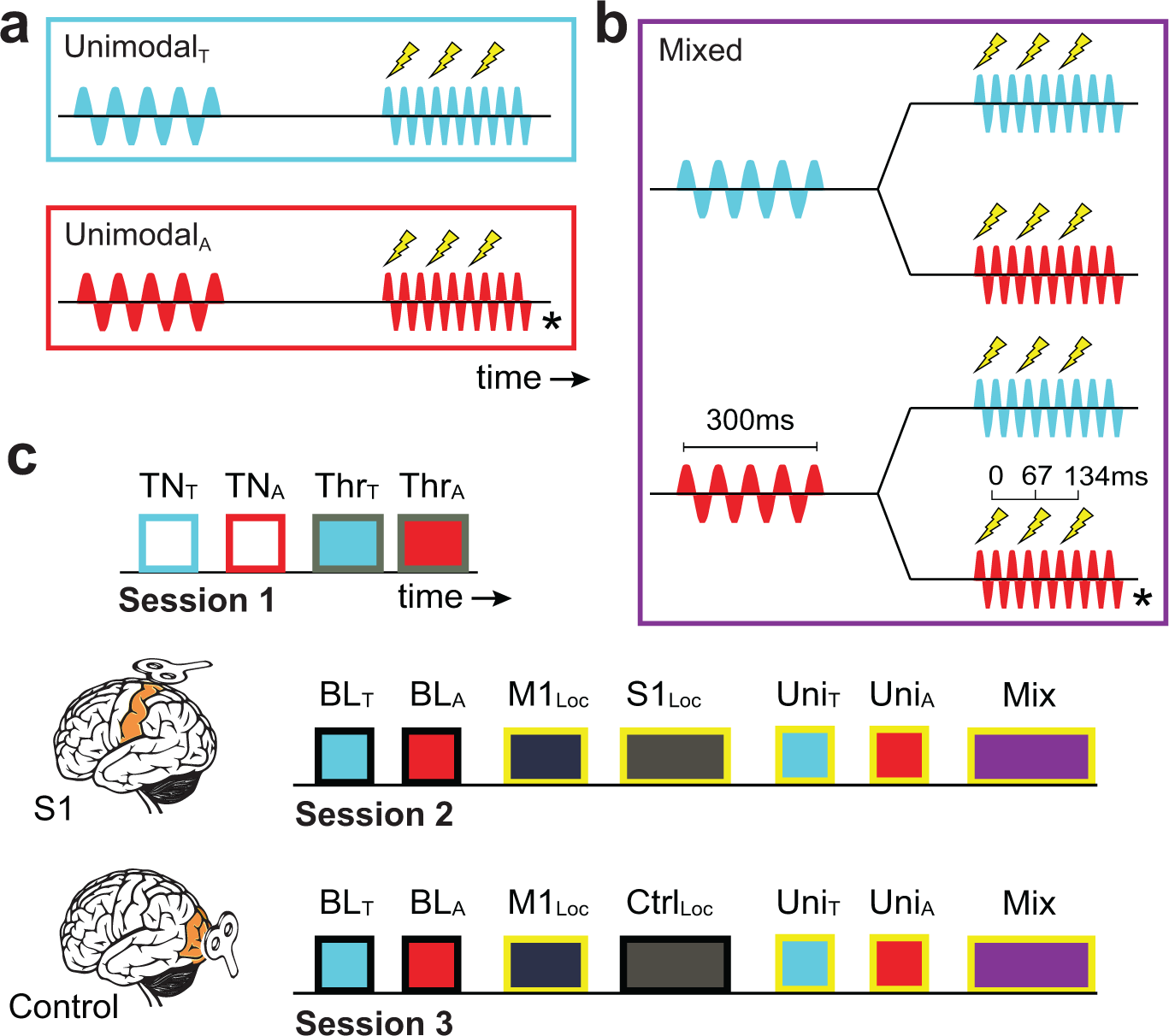
Experimental design. **a,** Frequency discrimination task in which the participant judged which stimulus was perceived to be higher in frequency on each trial. In blocks involving TMS, a burst (3 pulses) was delivered on each trial during the 2^nd^ interval. The Unimodal_T_ block (cyan box) comprised tactile stimuli only (cyan waveforms) and the Unimodal_A_ block (red box) comprised auditory stimuli only (red waveforms). In these blocks, subjects were required to attend to a single modality. **b**, The Mixed block (purple box) comprised trials requiring subjects to perform unimodal or crossmodal comparisons. In Mixed blocks, subjects were required to direct attention both modalities. **c**, The within-subjects design consisted of 3 sessions. Session 1 consisted of training blocks (TN_T_ and TN_A_) and blocks in which the subject’s tactile or auditory discrimination thresholds were measured (Thr_T_ and Thr_A_). Sessions 2 and 3 involved TMS (blocks with yellow outlines) over S1 or the control site. At the outset of each session, baseline blocks of auditory and tactile discrimination trials were completed without TMS (BL_T_ and BL_a_). The motor cortex hotspot was then localized (M1_Loc_) along with the target location over S1 (S1_Loc_) or the control site (Ctrl_Loc_). Subjects then performed the discrimination tasks in the Unimodal_T_, Unimodal_A_, and Mixed blocks involving TMS (Uni_T_, Uni_A_, Mix). The main analyses focused on the auditory-only trials denoted by the asterisks in *a* and *b* where the only factor that differed was the subject’s attention state.

To establish the TMS target site in each subject, we located the hand representation in left S1 and validated this target location using an independent perceptual task that has been shown to be sensitive to TMS (Methods). Prior to the discrimination task blocks, subjects performed a tactile localization task in which they reported on each trial whether they experienced a brief tap on the index finger of the left hand, right hand, both hands or no taps. In the absence of TMS, subjects achieved high performance (>0.90 probability correct) in all conditions (Fig. S1). Because S1 contains body representations that are predominantly contralateral, manipulating activity in left S1 using TMS should selectively impair the perception of taps on the right hand. Consistent with this prediction (Fig. 2a), online TMS delivered over left S1 significantly and systematically impaired performance on the localization task (F_3,42_= 17.6, p = 1.49e-07, η_p_^2^ = 0.55) by reducing performance accuracy on trials in which the tap was delivered to the right hand only (0.58 ± 0.09) and to both hands (0.39 ± 0.07). Critically, TMS-induced accuracy reductions resulted from a failure to detect taps on the right hand: Subjects reported feeling no stimulation when taps were delivered to the right hand only and they reported touch on only the left hand when taps were delivered to both hands (Fig. 2a). To quantify this lateralized TMS effect, we computed an extinction index (EI) for each subject where positive values indicate better performance on the hand ipsilateral to the stimulated cortex compared to the contralateral hand (Fig. 2b). The group-averaged EI (0.50 ±0.12) was significantly greater than 0 (t-test, t(14) = 4.19, p = 8.98e-04, d = 1.08) and positive EI values were observed in a significant number of individual subjects (13/15; binomial test, p = 0.003). These results demonstrate that TMS over S1 selectively disrupted the perception of simple taps on the hand contralateral to the stimulated hemisphere.

**Figure 2.**
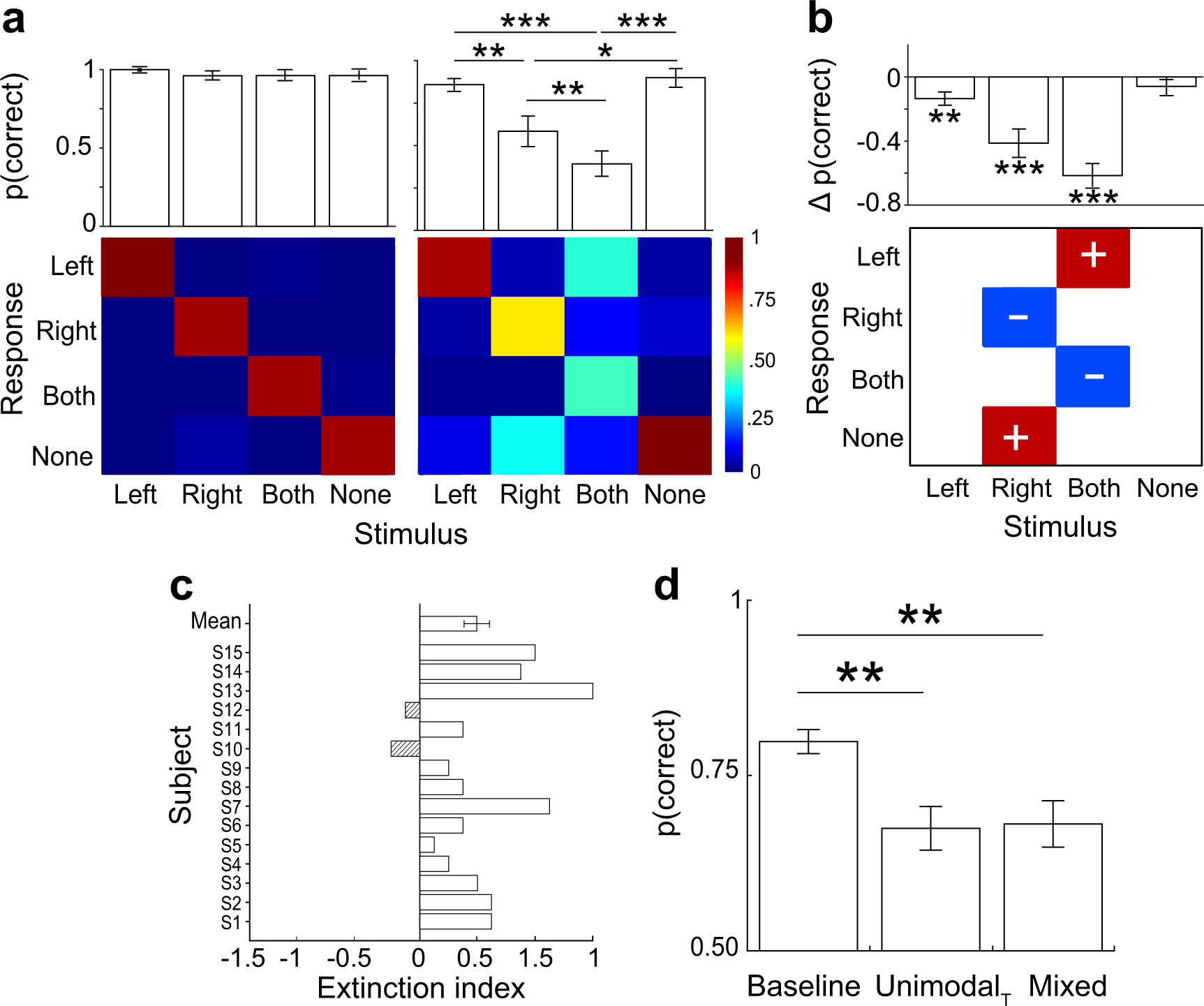
TMS over S1 selectively impairs contralateral touch. **a,** Group-averaged (n=15) tactile detection performance and response probabilities (confusion matrix) in participants detecting taps delivered to the index finger on the left hand, right hand, both hands, or no touch without TMS (left) and with TMS over left S1 (right). **b,** Impaired performance with TMS over left S1 is associated with reduced detection of taps on the right hand: Subjects are more likely to report no stimulation on ‘Right’ trials and touch on the left hand or no touch on ‘Both’ trials. Red and blue cells are conditions where response probabilities increased or decreased, respectively, by greater than 20% with TMS relative to performance without TMS. **c,** Individual and group-averaged extinction index (EI) values indicating relative TMS effects on detection performance on the left and right hands. Positive EI values indicate that TMS reduced detection performance on the right hand (contralateral to the stimulated cortex) more than the left hand. The two subjects who failed to show positive EI values were excluded from subsequent analyses. **d,** Tactile frequency discrimination performance (n=15) during Baseline, Unimodal_T_, and Mixed blocks. No TMS was applied during the Baseline block. Error bars indicate s.e.m. *** p<0.001, ** p<0.01, * p<0.05

In addition to impairing performance on the localization task, TMS over S1 also significantly impaired subjects’ ability to discriminate tactile frequency (Fig. 2c) (F_2,28_ = 11.25, p = 2.0e-04, η_p_^2^ = 0.45). In the absence of TMS, subjects successfully performed the tactile-only trials (0.80 ± 0.02). Relative to baseline performance, TMS over S1 significantly reduced performance in the Unimodal_T_ block (0.67 ± 0.03; t(14) = 3.76, p = 0.006, d = 0.97) and Mixed block (0.68 ± 0.03; t(14) = 3.74, p = 0.007, d = 0.97). Performance levels did not differ between the Unimodalj and Mixed blocks (t(14) = −0.28, p = 1, d = 0.07). TMS also delayed response times (RT) significantly (F_2,28_ = 20.0, p = 4.02e-06, η_p_^2^ = 0.59). Subjects took more time to respond in the Unimodal_T_ (796.1 ± 119.2msec; t(14) = −4.15, p = 0.003, d = 1.07) and Mixed blocks (898.8 ± 113.7msec; t(14) = −5.66, p = 1.75e-04, d = 1.46) compared to the baseline block (529.9 ±67.0msec). RT did not differ significantly between the Unimodal_T_ and Mixed blocks (t(14) = −2.03, p = 0.18, d = 0.52). These data indicate that manipulating S1 activity disrupted the ability of subjects to localize and discriminate tactile stimuli, thereby providing strong validation for the TMS target site over left S1.

To test the hypothesis that S1 becomes functionally coupled to cortical systems that mediate auditory frequency perception, we assessed subjects’ ability to discriminate sound frequency as we causally manipulated S1 activity under different attention states. If coupling between the somatosensory and auditory systems requires the deployment of attention to tactile frequency, we predicted that TMS over S1 would impair auditory discrimination performance only in the Mixed block during which subjects attended to both tactile and auditory frequency. We found that auditory frequency discrimination performance (Fig. 3a) differed significantly between the test blocks (F_2,28_ = 19.52, p = 4.9e-06, η_p_^2^ = 0.58). In the absence of TMS, subjects reliably and accurately performed the discrimination task (0.80 ± 0.02). Performance in the Unimodal_A_ block was nominally lower compared to baseline (0.77 ± 0.03), but this difference did not achieve statistical significance (t(14) = 2.01, p = 0.19, d = 0.52). This result implies that simply applying TMS over S1 was insufficient for inducing large changes in auditory frequency discrimination performance. Critically, performance on the auditory-only trials during the Mixed block (0.68 ± 0.03) was significantly impaired compared to performance in the baseline block (t(14) = 5.38, p = 2.9e-04, d = 1.39) and the Unimodal_A_ block (t(14) =4.37, p = 0.002, d = 1.13). For each subject, we computed a modulation index (defined as the performance difference between the Unimodal_A_ and Mixed blocks) and found significant accuracy reductions in the Mixed block (t(14) = 4.37, p = 6.32e-04, d = 1.13) that were ~10% on average across subjects (Fig. 3c). In fact, lower relative accuracies were observed in the Mixed block in nearly every subject (13/15; binomial test, p= 0.003). Importantly, in a control experiment that did not involve TMS (Fig. S2), no significant performance differences were observed between auditory-only trials in Unimodal_A_ and Mixed blocks (t(14) = 1.62, p = 0.13, d = 0.42; Unimodal_A_: 0.80 ± 0.01, Mixed: 0.77 ± 0.02). This control experiment result indicates that the observed decrements in auditory discrimination performance with TMS over S1 was not simply a consequence of requiring subjects to divide their attention over two senses. Additional Bayesian analyses (Methods) indicate substantially stronger evidence supporting the hypothesis that performance differs between the Unimodal_A_ and Mixed blocks with TMS over S1 (Bayes factor for S1-TMS: 57.43) compared to evidence in the control experiment without TMS (Bayes factor for No-TMS control experiment: 0.76). Thus, auditory frequency perception could be impaired only when TMS was applied over S1 as subjects attended to both tactile and auditory frequency.

**Figure 3.**
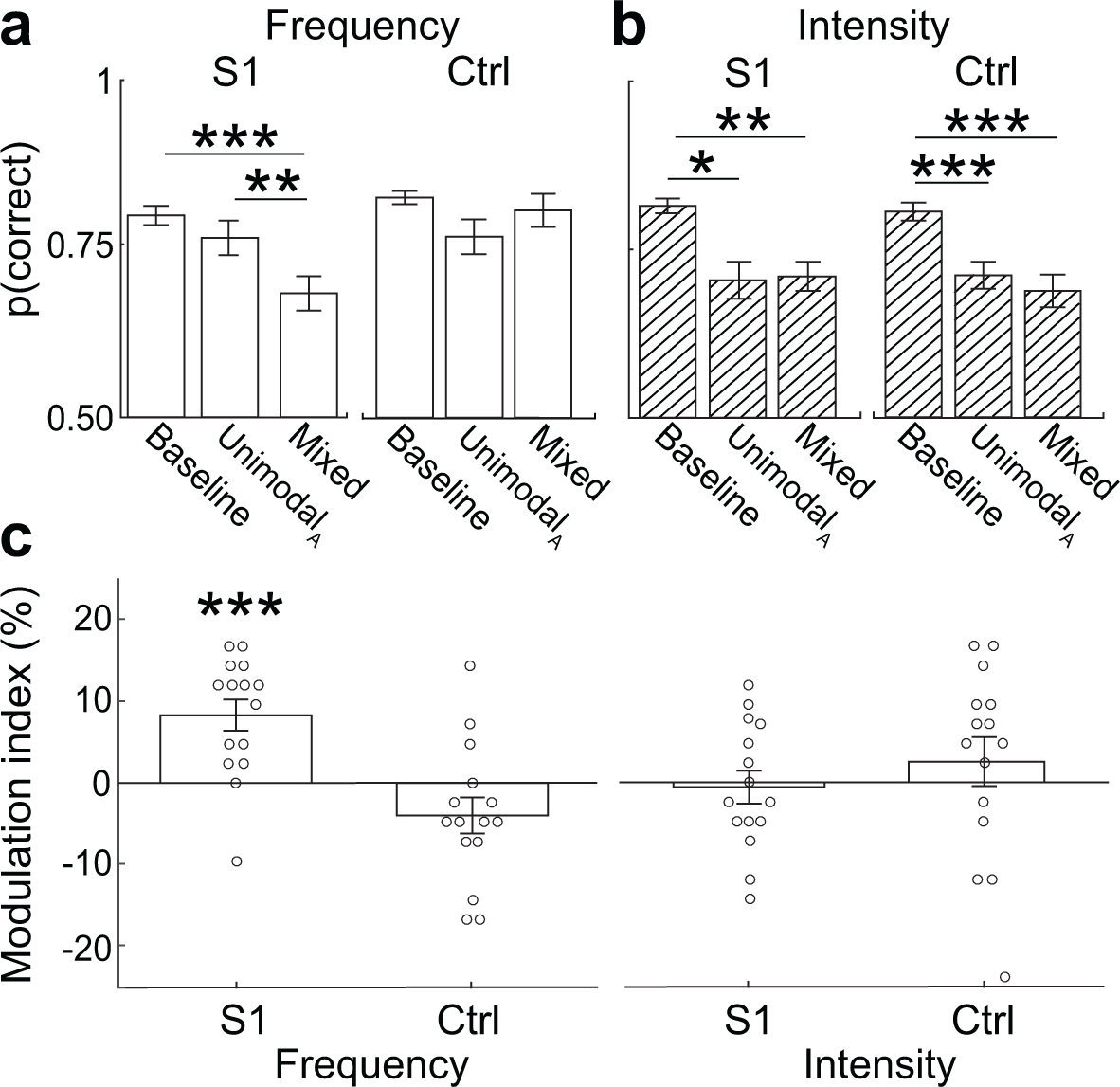
TMS over S1 impairs auditory perception. **a,** Performance on auditory-only frequency discrimination trials (n=15) for the different blocks (Baseline, Unimodal_A_, Mixed) and TMS sites. No TMS was applied during the Baseline block. Frequency perception is impaired only during the Mixed block by TMS over somatosensory cortex (S1), but not over a control site (Ctrl). **b,** Performance on auditory-only intensity discrimination trials (n=15). TMS impaired intensity perception regardless of block or site. **c,** Group-averaged (bar) and individual-subject (dots) modulation index (MI) values for frequency discrimination task (left) and intensity discrimination task (right) with TMS over S1 or over the control site indicating the absolute difference in performance accuracy between the Unimodal_A_ and Mixed blocks. Positive MI values indicate lower relative performance in the Mixed block. Error bars indicate s.e.m. *** p<0.001, ** p<0.01, * p<0.05.

TMS is associated with audible discharge sounds as well as tactile sensations on the scalp. These confounds can impact behaviour [21] and subjects may have been vulnerable to these non-specific TMS effects when they directed attention to auditory and tactile frequency. To address this possibility, we retested each subject while applying TMS over a control site (Methods). Auditory frequency discrimination performance with TMS over the control site differed significantly from performance achieved with TMS over S1 (Fig. 3a) (Site main effect: F_1,14_ = 10.98, p = 0.005,η_p_^2^ =0.44; Block main effect: F_2,28_ = 7.6, p = 0.002, η_p_^2^ = 0.35; Site*Block interaction: F_2,28_ = 12.43, p = 1.0e-04, η_p_^2^ = 0.47). These differences were due to the fact that TMS over the control site induced limited systematic changes in auditory discrimination performance (F_2,28_ = 3.01, p = 0.07, η_p_^2^ = 0.17; Baseline: 0.83 ± 0.01, UnimodalA: 0.77 ± 0.03, Mixed: 0.81 ± 0.02), in contrast to the robust patterned effects observed with TMS over S1. Although performance in the Unimodal_A_ block was nominally lower than baseline (t(14) = 2.33, p = 0.10, d = 0.60), no significant performance differences were observed in comparisons between the Mixed block and baseline (t(14) = 0.73, p = 1, d = 0.19) or the Unimodal_A_ block (t(14) = −1.81, p = 0.27, d = 0.47). Indeed, the data provided little support for the hypothesis that auditory frequency discrimination performance differed between the Unimodal_A_ and Mixed blocks with TMS over the control site (Bayes factor: 0.98). Thus, the performance pattern observed with TMS over S1 also cannot be attributed to non-specific TMS effects nor to an interaction between these effects and the attention manipulations. Notably, subjects (n = 15) who performed an analogous intensity discrimination task (Methods) were severely impaired by TMS regardless of stimulation site (Fig. 3b) (Site main effect: F_1,14_ = 0.28, p = 0.60, η_p_^2^ = 0.02; Block main effect: F_2,28_ = 21.7, p = 2.03e-06, η_p_^2^ = 0.6; Site*Block interaction: F_2,28_ = 0.47, p = 0.62, η_p_^2^ = 0.03). Specifically, auditory intensity discrimination performance differed significantly across blocks when TMS was delivered over both S1 (F_2,28_ = 1 1.76, p = 2.0e-04, η_p_^2^ = 0.45) and the control site (F228 = 13.96, p = 6.2e-05, η_p_^2^ = 0.50). Over both sites, TMS impaired performance in the Unimodal_A_ (S1: t(14) = 3.49, p = 0.01, d = 0.90; Ctrl: t(14) = 4.76, p = 8.97e-04, d = 1.23) and Mixed blocks (S1: t(14) = 4.37, p = 0.002, d = 1.13; Ctrl: t(14) = 5.12, p = 4.68e-04, d = 1.32). The lack of specificity in the TMS influences on intensity discrimination accuracy suggests that auditory intensity perception may be more vulnerable to non-specific TMS effects compared to auditory frequency perception. These results collectively establish the feature-dependence of TMS influences on audition and demonstrate that TMS *selectively* impaired auditory discrimination performance only when S1 activity was manipulated as subjects attended to auditory and tactile frequency.

Given the extensive connectivity within the sensorimotor cortical system, we considered whether impaired frequency discrimination performance resulted from TMS-induced excitability changes in motor cortex, which may have disrupted subjects’ ability to generate and report perceptual decisions correctly. To test this possibility, we compared EMG activity recorded from surface electrodes on the hand (Methods) during auditory-only trials in the Unimodal_A_ and Mixed blocks (Fig. S3). TMS over S1 rarely produced motor evoked potentials (MEPs) and similar rates of small-amplitude MEPs were measured in the two blocks (t(13) = −0.75, p = 0.47, d = 0.20) which did not differ in magnitude (t(16) = −0.43, p = 0.68, d = 0.20). Moreover, the amplitude of evoked responses was uncorrelated with task performance (r = −0.34, p = 0.17), ruling out the possibility that performance impairments in the Mixed block were due to TMS influences on motor cortex excitability.

According to the speed-accuracy tradeoff, auditory discrimination performance during the Mixed block could have been lower if subjects adopted different response criteria or performance strategies which caused them to rush their decisions. To test this possibility (Fig. S4), we evaluated response times with TMS over S1 (Baseline: 492.8 ± 47.1msec, Unimodal_A_: 748.7 ± 105.6msec, Mixed: 878.3 ± 89.7msec) and the control site (Baseline: 514.1 ± 44.9msec, Unimodal_A_: 831.8 ± 157.9msec, Mixed: 889 ± 122.5msec) and found significant RT variations over blocks (Block main effect: F_2,28_ = 7.51, p = 0.002, η_p_^2^ = 0.35), but no significant RT dependence on TMS site (Site main effect: F_1,14_ = 0.62, p = 0.44, η_p_^2^ = 0.04; Site*Block interaction: F_2,28_ = 0.18, p = 0.83, η_p_^2^ = 0.01). Critically, these results reveal that subjects took more time to respond when receiving TMS, although RT differences were only significant in comparisons between the Mixed and baseline blocks (S1: t(14) = −5.74, p = 1.53e-04, d = 1.48; Ctrl: t(14) = −4.30, p = 0.002, d = 1.11). Thus, the reduced discrimination accuracies in the Mixed block with TMS over S1 were not simply due to subjects hurrying their decisions. Moreover, the effect of TMS on RTs, which increases with attention load, potentially reflects a non-specific performance cost on auditory processing, as a similar pattern was also evident for the auditory intensity discrimination task (Fig. S4). This spatially-invariant, general TMS effect on the speed with which subjects respond clearly contrasts with the effects on frequency discrimination accuracy that are only obvious with TMS over S1.

Our main finding is that manipulating S1 activity using TMS impaired the ability to perform an auditory frequency discrimination task when subjects also attended to tactile frequency information. Manipulating attention alone or applying TMS over a control region did not alter performance accuracy significantly in the same frequency discrimination task. These behavioural results are consistent with the hypothesis that the functional coupling between the somatosensory and auditory systems is gated by directing attention to vibration frequency. We also found that TMS severely impaired auditory intensity perception regardless of attention state or TMS target. This pattern implies that TMS effects on audition differ depending on whether subjects perform a task that requires them to judge stimulus frequency rather than intensity, a result confirmed by a mixed-design ANOVA (Supplementary information), even when baseline performance on these tasks is standardized. Given the extensive perceptual interactions between touch and audition in the temporal frequency domain [10,18,22], it is perhaps unsurprising that neural activity in S1 can influence auditory processing – our data suggest that even TMS-evoked neural signals in S1, in the absence of bottom-up stimulus-driven activity, may be transmitted to neural circuits that support audition. TMS-evoked S1 activity could propagate along a number of cortico-cortical and thalamocortical pathways which connect the somatosensory and auditory systems [23–25]. Presumably, neural activity is similarly routed through these networks when actual sensory inputs are processed, which result in the co-activation of the somatosensory and auditory cortical systems observed in earlier studies [4,9,26]. Outside of the sensory cortices, auditory and tactile information can also be represented in common or overlapping neural populations in frontal regions that support working memory and decision-making [27], which may have been remotely modulated by TMS over S1.

A major limitation of our study is that we infer how attention and TMS engage perceptual systems based solely on behavioural data. Our results would support a critical role for feature-based attention in dynamically regulating the network coupling, assuming that the behavioural effects observed in our subjects truly reflect remote TMS effects over a distributed frequency processing network that supports audition and touch. This assumption must be tested in future experiments that explicitly monitor brain activity and characterize network architecture. It should be noted that our data is also consistent with alternative accounts that do not assume remote TMS effects. Although we targeted somatosensory cortex, it is possible that we delivered TMS over a more posterior region in parietal cortex (Methods). Because posterior parietal cortex (PPC) has been shown to support working memory [28], perceptual decision making [29], attentional selection [30], and representations of auditory frequency information [8], impaired performance on the auditory frequency discrimination task may have resulted from local TMS effects in PPC. In each of these cases, though, the effect of TMS on auditory frequency perception would still be conditioned on attention state, so our primary finding that attention to both touch and audition is required for TMS to impair auditory frequency perception applies to accounts assuming either local or remote effects. We favour the latter, in part, based on results of combined brain stimulation with human neuroimaging studies that demonstrated how attention state modulates the coupling between fronto-parietal control networks and sensory cortex [31,32]. Our results potentially build on these earlier findings by showing that TMS may also reveal state-dependent coupling, as indexed by highly selective behavior changes, between cortical systems that are thought to support different sensory modalities which process the same stimulus features.

How might selective attention gate the crossmodal coupling between the somatosensory and auditory systems? A prevailing view is that oscillatory dynamics in intrinsic brain activity control the flow of information through anatomical pathways [33,34]. By modulating the effective connectivity between different neural populations and networks, such mechanisms are hypothesized to support flexible and context-dependent behaviours. Selective attention may modulate neuronal oscillations [35]: Increasing the coherence between oscillatory activity in different neuronal populations can facilitate information transmission and integration [36–38]. In our paradigm, directing attention to tactile frequency may have synchronized the intrinsic activity in S1 and the cortical networks that support frequency processing for touch and audition. This coherent network state would then enable the perturbations induced by TMS over S1 to propagate to neural populations that mediate auditory frequency processing thereby disrupting auditory perception. If similar state-dependent transmission of stimulus-evoked activity underlies the perceptual interactions between audition and touch, our results could be interpreted as evidence for the general framework that multisensory interactions result from crossmodal binding through neural coherence [39].

Sensory cortical systems that are traditionally considered to be dedicated to individual modalities often co-activate, even when the inputs are presented in a single modality [16]. These co-activations are thought to reflect the binding of information-linked neural representations via the crossmodal spread of spatial attention [40,41] or feature-based attention [42–44]. The causal manipulation of activity in one sensory system using noninvasive brain stimulation can modulate processing in a different sensory modality [45], implying an interactive connectivity between sensory cortical systems. Our results demonstrate that the interactive coupling between cortical systems which support different sensory modalities is modulated by selective attention. This gating of crossmodal coupling by selective attention may critically support multisensory interactions and feature-specific perception.

## METHODS

### Participants

A total of 37 subjects participated in this study. Fifteen subjects (9 males; mean age ± SD: 23.73 ± 6.49 years) participated in the frequency discrimination experiment (Experiment 1). Fifteen subjects (7 males, mean age ± SD: 25.2 ± 6.86 years) participated in the intensity discrimination experiment (Experiment 2). Fifteen subjects (5 males; mean age ± SD: 26.2 ± 4.9 years) participated in the frequency discrimination experiment without TMS (Experiment 3). Two subjects participated in both Experiments 1 and 2. Two subjects took part in all three experiments. Two subjects participated in both Experiments 1 and 3. Six subjects were left handed, according to the Edinburgh Handedness Inventory [46]. All participants reported normal tactile and auditory sensibilities. No participant reported a neurological or psychiatric history. No subjects reported contraindications to non-invasive transcranial brain stimulation [47]. All testing procedures were performed in compliance with the policies and procedures of the Baylor College of Medicine Institutional Review Board. All participants gave their written informed consent and were paid for their participation.

### Stimuli and procedures

#### Stimuli

Auditory and tactile stimuli tested in the discrimination experiments comprised sine waves (sample rate: 44.1kHz; linear ramp: 30msec) that were digitally generated in Matlab (2011b, MathWorks) and presented with Psychtoolbox-3 [48] running on a MacBook Pro (model A1278; OS × 10.9.5, 2.5 GHz Core i5, 4 GB of RAM). Auditory stimuli consisted of analog signals from one channel of the auxiliary port which were amplified (PTA2, Pyle) and delivered binaurally via noise-cancelling in-ear headphones (ATH-ANC23, Audio-Technica U.S., Inc). Participants also wore noise-attenuating earmuffs (Peltor H10A Optime 105 Earmuff, 3M) over the in-ear headphones which served to attenuate the sounds associated with TMS discharge and tactile stimulation. Tactile stimuli consisted of analog signals from the other channel of the auxiliary port which were amplified (Krohn-Hite Wideband Power Amplifier, model 7500) and delivered to the subject’s right index finger through an electromechanical tactor (type C-2, Engineering Acoustics, Inc.). The stimuli tested in the tactile localization task consisted of brief taps delivered through a pair of miniature electromechanical tactors (type C-FT, Engineering Acoustics, Inc.) attached to the left and right index fingers. Tactors were fastened to the distal phalanges using self-adherent cohesive wrap bandages.

#### General procedures

The frequency and intensity discrimination experiments (Experiments 1 and 2, respectively) each comprised 3 sessions. Each session took place on separate days (mean inter-session interval ± SD, Experiment 1: 4.5 ± 2.7 days; Experiment 2: 3.8 ± 2.2). The general organization of the sessions was the same for the frequency and intensity discrimination experiments and the general procedures are detailed using the frequency discrimination experiment as the example (Fig. 1).

During session 1, participants were trained to perform the frequency discrimination task in both the auditory and the tactile modalities. After the initial training period, each participant’s auditory and tactile frequency discrimination thresholds (FDT_A_ and FDT_T_) were estimated with a Bayesian adaptive threshold-tracking procedure [49] (Supplementary information). We defined FDT as the minimum change in stimulus frequency required for a comparison stimulus to be perceived as higher in frequency compared to a 200-Hz standard stimulus 80% of the time. For the intensity discrimination experiment, we estimated each participant’s intensity discrimination threshold (IDT), defined as the minimum change in stimulus amplitude required for a 200-Hz comparison stimulus to be perceived as more intense than a 200-Hz reference stimulus of a fixed supra-threshold intensity 80% of the time (Supplementary information). In preliminary experiments, we identified reference amplitudes for 200-Hz auditory and tactile stimuli such that they were perceived as equally intense (Supplementary information) in order to perform the crossmodal intensity judgments (Fig. S5). In sessions 2 and 3, subjects were tested with stimuli determined according to their FDT or IDT (see below), which enabled us to standardize baseline performance across subjects and tasks.

Sessions 2 and 3 involved TMS and were identical except for the location over which TMS was applied. The order of the sessions involving TMS over the S1 site or the control site was counterbalanced across subjects. Each session began with 2 baseline blocks during which subjects performed auditory-only or tactile-only frequency discrimination trials. Performance on these blocks, which were achieved in the absence of TMS, established the consistency of FDT values estimated in session 1 and provided a baseline against which we compared discrimination performance achieved with TMS. After the subject completed the baseline blocks, we performed TMS mapping to localize the motor hotspot in the subject’s left motor cortex and to estimate her resting motor threshold (RMT) (see *M1 hotspot localization and RMT estimation*). After establishing RMT, the S1 site or control site was localized (see *Localization of S1 and control sites*) and participants began behavioral testing with TMS. In sessions with TMS over S1, subjects performed a tactile localization task before the discrimination task. For the discrimination task, each participant was tested on 1 Unimodal_A_ block (auditory-only trials), 1 Unimodal-Γ block (tactile-only trials), and 3 Mixed blocks (auditory-only, tactile-only, and crossmodal trials). Each block contained 42 trials and block order was randomized in each subject. Subjects were provided with 5-min rest intervals between each block.

#### Tactile localization task

Subjects performed a 4-alternative forced choice (4AFC) tactile localization task with and without TMS. We included this task as an independent method for validating TMS coil position over S1 given that similar paradigms have been used previously to assess TMS effects on touch [50–52] and their dependence on task demands [53]. On each trial, a brief (5-msec) tactile tap was delivered to the index finger on either the left hand, right hand, both hands, or there was no stimulation. Subjects, who maintained fixation throughout the trials, received a visual cue indicating the trial interval during which the tap(s) could have been delivered followed by a cue indicating that the subject should respond. On each trial, subjects verbally reported whether they perceived touch on the left hand only (“Left”), right hand only (“Right”), both hands simultaneously (“Both”), or no stimulation (“None”). Tactile-stimulation trials (left, right, and both) were repeated 8 times each and no-stimulation trials were repeated 4 times in random order. The amplitude of the taps was set at 120% of each subject’s tactile detection thresholds, determined with an adaptive threshold tracking procedure [49] (Supplementary information). These amplitudes were chosen to standardize performance across subjects at ~90% in the absence of TMS (Fig. S1).

#### Frequency discrimination task

Participants discriminated auditory and tactile frequencies in a 2-interval, 2-alternative forced choice (2AFC) paradigm (stimulus duration: 300msec; inter-stimulus interval: 500msec). Subjects were asked to report which interval contained the stimulus perceived to be higher in frequency by button press using their left hand. Throughout the test blocks, subjects maintained their gaze on a central fixation point on a computer screen.

Trials were organized according to 3 modality conditions (auditory-only, tactile-only, crossmodal). Within each modality condition, there were 3 frequency conditions resulting in a total of 9 unique stimulus pairs in Experiment 1 (Table S1). During each unimodal block, subjects were tested on 3 stimulus pairs (all of the same modality condition) individualized according to each subject’s FDT estimates: 200Hz vs 200Hz-FDT, 200Hz vs 200Hz+FDT, and 200Hz+FDT vs 200Hz+2*FDT. The average (±SD) FDT_A_ and FDT_T_ were 3 ± 2Hz and 39 ± 16Hz, respectively. These thresholds, relative to 200Hz, are consistent with published reports [10,11]. Each stimulus pair was repeated 14 times in the unimodal blocks. During the Mixed blocks, the auditory-only and tactile-only trials were interleaved with crossmodal trials that also comprised 3 stimulus pairs individualized according to FDT: (auditory frequency vs tactile frequency) 200Hz vs 200Hz-FDT_T_, 200Hz vs 200Hz+FDT_T_, and 200Hz+2*FDT_T_ vs 200Hz+FDT_T_. These crossmodal trials required subjects to compare the frequency of a sound to the frequency of a vibration within a single trial [27]. Because tactile frequency discrimination thresholds are substantially larger than auditory frequency discrimination thresholds [11], the limiting factor in participants’ ability to perform crossmodal frequency comparisons is their tactile sensitivity. Accordingly, we defined the crossmodal frequency pairs for each subject with respect to his or her FDT_T_ (Table S1). This ensured that subjects were able to perform the crossmodal frequency comparisons (Fig. S5) at levels exceeding chance level even while receiving TMS, irrespective of TMS site (t(14) = −0.41, p = 0.68, d =0.11; S1 session: 0.71 ± 0.02; Ctrl session: 0.72 ± 0.03). Over 3 Mixed blocks (42 trials each), each of the 9 unimodal and crossmodal stimulus pairs were repeated 14 times. During all blocks, the stimuli comprising each stimulus pair were presented in random order on each trial and the ordering of stimulus pairs was randomized over the block.

We took multiple steps to ensure that participants performed the frequency discrimination task using frequency rather than intensity cues [10–13]. First, the stimuli were equated for perceived intensity in preliminary experiments (Supplementary information). Additionally, a random jitter (±10%) was applied to the subjectively-matched amplitudes on each trial to guarantee that differences in perceived intensity did not covary with frequency differences.

#### Intensity discrimination task

Participants discriminated auditory and tactile intensities in a 2AFC paradigm analogous to the frequency discrimination task. On each trial, subjects reported which of two supra-threshold stimuli was perceived to be more intense (stimulus duration: 300msec; inter-stimulus interval: 500msec; stimulus frequency: 200Hz). As in the frequency discrimination experiment, each subject was tested on 9 unique stimulus pairs (Table S2). To standardize baseline performance to ~80%, stimulus amplitudes were determined according to IDT_A_ and IDT_T_ established with adaptive threshold tracking procedures (Supplementary information). The average (±SD) IDT_A_ and IDT_T_ were 16.5 ± 9.8% of the auditory reference amplitude and 10.5 ± 4.1% of the tactile reference amplitude, respectively. The reference auditory and tactile signals measured as the output of the amplifiers were 1.9V and 2.3V, respectively.

### Transcranial Magnetic Stimulation

#### TMS setup and stimulation parameters

Bi-phasic pulses were delivered through a figure-of-eight coil (D70^2^ coil, wing diameter: 70mm) connected to a Magstim Rapid^2^ stimulator unit (The Magstim Company). The location of the TMS coil over the participant’s scalp was continuously tracked during the experiment using a frameless stereotaxic neuronavigation system (Brainsight, version 2.3.1, Rogue Research, Inc). Note that we did not perform neuronavigation based on each subject’s anatomy; however, we instead used the Brainsight system to monitor the relative positions of the TMS coil and each participant’s scalp (in an arbitrary coordinate system defined by a common anatomical brain scan referenced to all subjects) in order to landmark TMS targets, to verify accurate TMS targeting online, and to perform electromyography recordings (see below).

During discrimination trials in which TMS was applied, a train of 3 pulses (inter-pulse interval: 67ms) was delivered over the TMS target site at the start of the second stimulus interval (Fig. 1a,b). TMS intensity was determined according to each subject’s RMT. Although we aimed to stimulate all subjects at 120% RMT, we reduced TMS intensity to 110% RMT in subjects (6 in Experiment 1 and 5 in Experiment 2) who reported discomfort at 120% RMT. TMS intensity, expressed as a percentage of maximum stimulator output (MSO), was 61.8 ± 5% (Experiment 1) and 65.3 ± 7.13% (Experiment 2). TMS timing was controlled by Matlab and TMS trains were triggered by TTL pulses sent via a DAQ device (model USB-1208FS, Measurement Computing Corporation).

During trials in the detection task in which TMS was applied, a pair of TMS pulses (interpulse interval: 40ms) was delivered over S1. Relative to the time of the tactile stimulus, the first TMS pulse preceded the tap by 10msec [54,55]. TMS pulses were presented at the same relative time on trials that did not include tactile stimulation.

#### Electromyography (EMG)

We recorded EMG activity from the first dorsal interosseous (FDI) muscle on the right hand using the built-in EMG setup in the Brainsight system. Two pre-gelled, disposable Ag/AgCl surface electrodes (Kendall Medi-Trace mini electrodes) were positioned over the FDI muscle while a ground electrode was placed over the styloid process of the ulna bone. Before placing the EMG electrodes, the skin was scrubbed with an alcohol wipe to reduce impedance.

#### M1 hotspot localization and RMT estimation

For each participant, we first localized the motor hotspot, the scalp site over left motor cortex (M1) where we reliably produced motor evoked potentials (MEPs) in the right FDI with a single TMS pulse using standard methods [56]. After localizing the motor hotspot, we established the subject’s RMT, defined as the lowest stimulation intensity that evoked a response in the relaxed FDI muscle of 100-μν peak-to-peak amplitude in 5 out of 10 trials. The average RMT was 53 ± 4% MSO (Experiment 1) and 56 ± 7% MSO (Experiment 2). The localization of the M1 hotspot and estimation of RMT was performed in sessions 2 and 3 and RMT values were highly consistent across days (Fig. S6).

### Localization of S1 and control sites

To localize the S1 site, we positioned the TMS coil over the M1 hotspot and applied single TMS pulses at 120% RMT as we systematically moved the coil posteriorly in 5-mm increments. The S1 site was the first location in which no MEPs were produced and subjects reported no TMS-associated sensations of muscle activity. On average, this S1 site was posterior to the M1 hotspot by 28 ± 8mm (Experiment 1) and 27 ± 7mm (Experiment 2). These distances, while offering no definitive assurance that S1 was targeted, fall within the range of analogous distances reported in previous studies presuming TMS over S1 [51,53,57–60]. For TMS over the control site, the TMS coil was positioned 3cm above the inion on the midline with the handle pointing upward.

### Data analysis

Statistical analyses were performed in Matlab (2011b) and R (R Studio version 0.99.892; Bayes Factor package). We performed normality tests on all data using Kolmogorov-Smirnov tests.

#### Tactile localization task

Our primary analysis of the localization task was aimed at characterizing lateralized effects of TMS on tactile detection in the 4AFC task.

Accordingly, we defined an extinction index (FI) as:

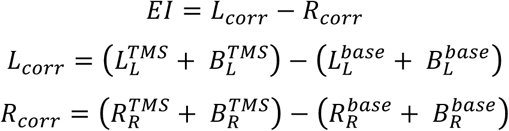

where *L_corr_* and *R_corr_* corresponded to baseline-corrected detection rates on the left and right hands, respectively. Baseline-corrected rates were calculated as the difference in the detection rates achieved with TMS and without TMS (base). For the TMS and base-conditions, detection rates on each hand were summed over unimanual (L or R) and bimanual (B) trials. Positive EI values indicate higher baseline-corrected detection rates on the left hand (ipsilateral to the S1 cortex receiving TMS) compared to the right hand (contralateral to the S1 cortex receiving TMS). Because S1 contains a predominantly contralateral hand representation, we predicted TMS should selectively impair detection on the right hand if TMS disrupted perception at all [54,55].

#### Frequency and intensity discrimination tasks

Performance accuracy was quantified for each modality condition within the Baseline, Unimodal, and Mixed blocks. Because frequency discrimination performance did not differ significantly across stimulus pairs (Fig. S7), we collapsed over the frequency condition in the main analyses. For instance, performance on auditory-only trials was computed over 42 trials in the Baseline, Unimodal_A_, and Mixed blocks. We similarly collapsed over the amplitude condition in analyses of the intensity discrimination task (Fig. S8).

In group-level analysis, we first performed a two-way repeated-measures ANOVA with Block (Baseline, Unimodal, Mixed) and Site (S1 and control) as within-subjects factors to test whether TMS effects on auditory-only trials differed according to the attention manipulation and the site of stimulation (Fig. 3). We then conducted separate one-way repeated-measures ANOVAs on data from the S1 and control site sessions with Block as the within-subjects factor followed by post hoc tests which were adjusted for multiple comparisons using the Bonferroni correction (corrected p-values are reported in the text). Data from the intensity discrimination task in Experiment 2 were similarly analyzed. To summarize succinctly the modulatory effect of the attention manipulation on task performance, we calculated a modulation index (MI) for each task and site as the accuracy difference between the Mixed and Unimodal blocks (Unimodal minus Mixed). Positive MI values indicate a relative impairment in accuracy during the Mixed block. Using one-sample t-tests, we tested the hypothesis that MI values were significantly different from 0 (Fig. 3c). We additionally conducted Bayes factor analyses to quantify the relative support for the hypothesis that performance differed in the Unimodal and Mixed blocks (H_1_) compared to the null hypothesis of no Unimodal and Mixed block performance differences (H_0_) where the Bayes factor was p(data|H_1_)/ p(data|H_0_).

#### EMG trace analysis

EMG data were analyzed for subjects included in the group-level analyses of the discrimination data. (Note that EMG data were available for 14/15 subjects in Experiment 1 due to a technical problem.) Trial-wise EMG traces were sorted according to modality condition (auditory-only, tactile-only trials) and block (Unimodal_A_, Unimodal_T_, Mixed). EMG traces were visually inspected to identify trials containing MEPs. For each condition and block, we calculated the likelihood that a TMS pulse produced a MEP. In group-level tests, we compared these likelihoods for the unimodal and the Mixed blocks using 2-tailed paired t-tests for each modality condition separately. To calculate MEP amplitudes, we computed the difference between the maximum and minimum voltages recorded in the interval ranging from 19–65msec after each TMS pulse. Restricting our analyses to this interval served to exclude artefactual voltage changes related to TMS discharge. In group-level tests, we compared MEP amplitudes between the unimodal and Mixed blocks for each modality condition using a two-sample t-test.

## Acknowledgements

This work was performed in the Neuromodulation and Behavioral Testing Facilities of BCM’s Core for Advanced MRI (CAMRI). This work was supported by the National Institutes of Health (R01NS097462) and the Alfred P. Sloan Research Fellowship. We thank members of the Yau lab for helpful discussions.

